# *Distal-less* homeobox genes *Dlx5/6* regulate Müllerian duct regression

**DOI:** 10.1101/2022.04.08.487506

**Authors:** Rachel D. Mullen, Brice Bellessort, Giovanni Levi, Richard R. Behringer

**Affiliations:** Department of Genetics, University of Texas MD Anderson Cancer Center, Houston, Texas, U.S.A; Physiologie Moléculaire et Adaptation, CNRS UMR7221, Muséum National d’Histoire Naturelle, Département AVIV, Paris, France

**Keywords:** sex differentiation, anti-Müllerian hormone, reproductive tract development, *Dlx5*, *Dlx6*, *Amhr2*

## Abstract

*Dlx5* and *Dlx6* encode distal-less homeodomain transcription factors that are present in the genome as a linked pair at a single locus. *Dlx5* and *Dlx6* have redundant roles in craniofacial, skeletal, and uterine development. Previously, we performed a transcriptome comparison for anti-Müllerian hormone (AMH)-induced genes expressed in the Müllerian duct mesenchyme of male and female mouse embryos. In that study, we found that *Dlx5* transcripts were nearly seven-fold higher in males compared to females and *Dlx6* transcripts were found only in males, suggesting they may be AMH-induced genes. Therefore, we investigated the role of *Dlx5* and *Dlx6* during AMH-induced Müllerian duct regression. We found that *Dlx5* was detected in the male Müllerian duct mesenchyme from E14.5 to E16.5. In contrast, in female embryos *Dlx5* was detected in the Müllerian duct epithelium. *Dlx6* expression in Müllerian duct mesenchyme was restricted to males. *Dlx6* expression was not detected in female Müllerian duct mesenchyme or epithelium. Genetic experiments showed that AMH signaling is necessary for *Dlx5* and *Dlx6* expression. Müllerian duct regression was variable in *Dlx5* homozygous mutant males at E16.5, ranging from regression like controls to a block in Müllerian duct regression. In E16.5 *Dlx6* homozygous mutants, Müllerian duct tissue persisted primarily in the region adjacent to the testes. In *Dlx5-6* double homozygous mutant males Müllerian duct regression was also found to be incomplete but more severe than either single mutant. These studies suggest that *Dlx5* and *Dlx6* act redundantly to mediate AMH-induced Müllerian duct regression during male differentiation.

## Introduction

The male reproductive tract organs include the vas deferentia, epididymides, and seminal vesicles. These structures provide the conduit for movement and maturation of spermatozoa from the testes for sexual reproduction. While formation and differentiation of these male reproductive tract organs are essential for reproduction, another process which eliminates a progenitor organ system termed Müllerian duct regression is also required for male development (1). These developmental processes are regulated by the presence or absence of fetal hormones (2).

The reproductive tract organs of mammals are derived from two pairs of epithelial tubes surrounded by mesenchyme called the Wolffian ducts (progenitor male reproductive tract) and Müllerian ducts (progenitor female reproductive tract). The Wolffian ducts differentiate into the vas deferentia, epididymides, and seminal vesicles. The oviducts, uterus and upper vagina are derived from the Müllerian ducts. Interestingly, both the Wolffian and Müllerian ducts are formed regardless of the genetic sex of the embryo. Differential hormone signaling after the formation of these genital ducts results in Wolffian duct loss in female embryos and Müllerian duct regression in male embryos.

Müllerian duct regression is mediated by the TGF-β family member anti-Müllerian hormone (AMH) secreted by Sertoli cells of the fetal testes (3). AMH signaling acts in the Müllerian duct mesenchyme through ACVR1 or BMPR1A type 1 receptors (shared with the bone morphogenetic protein (BMP) signaling pathway) and a sole anti-Müllerian hormone type 2 receptor (AMHR2) (4,5). *Amhr2* is expressed in the Müllerian duct mesenchyme and in the somatic cells of the gonads (6). In the Mullerian duct mesenchyme, AMH binds AMHR2 which then activates its type 1 receptors. This AMH ligand receptor complex subsequently phosphorylates R-SMAD1, 5 and 8 (also shared with the BMP signaling pathway) (4). These redundant phosphorylated R-SMADs then presumably activate downstream effectors of AMH signaling.

*Amh* and *Amhr2* are each required for Müllerian duct regression. Male mice homozygous for either *Amh* or *Amhr2* null alleles retain Müllerian duct derived structures, including a uterus and oviducts (7,8). Further, mutations of either the *AMH* or *AMHR2* genes in human males result in Persistent Mullerian Duct Syndrome (PMDS). Like mouse models with *Amh* and *Amhr2* loss, patients with PMDS have testes and male secondary sex characteristics but retain female reproductive tract organs associated with their male reproductive tract (9–11). A mutation in *Amhr2* in dogs is also known to result in PMDS (12). Males of these species with loss of either *AMH* or *AMHR2* also have subfertility likely due to a combination of the effects of cryptorchidism (observed in humans and dogs) and the presence of superimposed Müllerian duct tissues impeding sperm passage (mouse, human and dog) (9,12). *AMH* alone is also sufficient for Müllerian duct regression in female embryos. Transgenic female mice with widespread expression of human AMH have complete regression of the Müllerian ducts and lack a uterus and oviducts (13).

Few downstream effectors of AMH have been identified. In mice, loss of the signaling molecule beta-catenin encoded by *Ctnnb1* in the Müllerian duct mesenchyme results in the complete retention of the Müllerian duct in newborn males (14). In a recent RNA-seq study we compared male and female mouse Müllerian duct mesenchyme transcriptomes soon after the initiation of AMH signaling in males to identify potential AMH-induced target genes. In that study we identified the BMP target *Osterix/Sp7 (Osx)* (15,16). We found that AMH signaling is necessary and sufficient for *Osx* expression. *Osx* null male mice have a delay in Müllerian duct regression but later Müllerian duct derived tissues are eliminated (16).

This transcriptome analysis also revealed differential upregulation of BMP target genes *Dlx5* and *Dlx6* in the Müllerian mesenchyme of males in comparison to females. Based on this transcriptome data and that the BMP pathway shares multiple signaling components with the AMH signaling pathway, we further characterize *Dlx5* and *Dlx6* expression and function during Müllerian regression. We show that AMH signaling is necessary for *Dlx5* and *Dlx6* expression in Müllerian duct mesenchyme. Loss of *Dlx5* or *Dlx6* results in partial retention of the Müllerian ducts in male mice, whereas double mutant males consistently retain more Müllerian tissue. This study identifies *Dlx5* and *Dlx6* as redundant downstream effectors of AMH signaling for Müllerian regression.

## Materials and Methods

### Mice

Swiss outbred mice were obtained from Taconic Biosciences. C57BL/6J mice were obtained from the Jackson Laboratory. *Dlx6^tm1Jlr^* mice were obtained from the MMRRC (Mutant Mouse Resource and Research Center). *Amhr2^tm3(cre)Bhr^ (Amhr2-Cre,* (17)), *Amhr2^tm2Bhr^ (Amhr2-lacZ,* (18)), *Dlx5^tm1Levi^ (Dlx5-lacZ,* (19))*, Dlx6^tm1Jlr^ (Dlx6-lacZ,* (20)), *Dlx5/Dlx6^tm1Levi^* (*Dlx5-6*, (21)) mice were maintained on a predominantly C57BL/6J genetic background. *Amhr2-Cre, Amhr2-lacZ, Dlx5-lacZ, Dlx6-lacZ,* and *Dlx5-6* mice were genotyped by PCR as described previously. All animal procedures were approved by the Institutional Animal Care and Use Committee of the University of Texas MD Anderson Cancer Center. Studies were performed consistent with the National Institutes of Health Guide for the Care and Use of Laboratory Animals.

### β-galactosidase staining

*Dlx5-lacZ* and *Dlx6-lacZ* heterozygous males were bred with Swiss females to establish timed matings for β-galactosidase (β-gal) staining as previously described (16).

### Immunofluorescence

Immunofluorescence was performed as previously described (22). Rabbit anti-DLX5 polyclonal antibody (Sigma-Aldrich Cat# HPA005670, RRID:AB_1078681, 1:50). Rabbit anti-PAX2 polyclonal antibody (Thermo Fisher Scientific Cat# 71-6000, RRID:AB_2533990, 1:100). Primary antibodies were detected with goat anti-rabbit IgG (H+L) Alexa Fluor Plus 488 rabbit (Thermo Fisher Scientific Cat# A32731TR, RRID:AB_2866491, 1:200). At least three specimens of each genotype were analyzed.

## Results

### *Dlx5* and *Dlx6* expression in the developing reproductive organs is sexually dimorphic

Previously, we generated transcriptomes by RNA-seq of FACS-purified Müllerian duct mesenchyme cells from E14.5 male and female embryos to identify candidate genes induced by AMH that mediate Müllerian duct regression (16). At this stage of development, male mesenchymal cells are responding to AMH secreted by the testes, whereas in female embryos these cells are naïve to AMH because the fetal ovaries do not express AMH. Analysis of the bulk RNA-seq results showed that *Dlx5* transcripts were 6.7-fold higher in males compared to females and *Dlx6* transcripts were found only in males, suggesting these genes may respond to AMH signaling to mediate Müllerian duct regression.

Previous studies in the mouse showed that *Dlx5* is expressed in the epithelium of the E18.5 uterus and subsequently in the luminal and glandular epithelia of the postnatal and adult uterus (23). In that study, expression was also detected earlier in the Müllerian ducts at E15.5 and E16.5. We examined DLX5 expression in the Müllerian ducts of E14.5 male and female embryos by immunofluorescence (**Fig. 1**). In males, DLX5 was detected in the mesenchymal cells surrounding the Müllerian duct epithelium (**Fig. 1A**). In contrast, but consistent with previous studies (23), DLX5 expression was detected in the epithelial cells of the Müllerian duct of E14.5 female embryos (**Fig. 1B**).

**Fig. 1.**
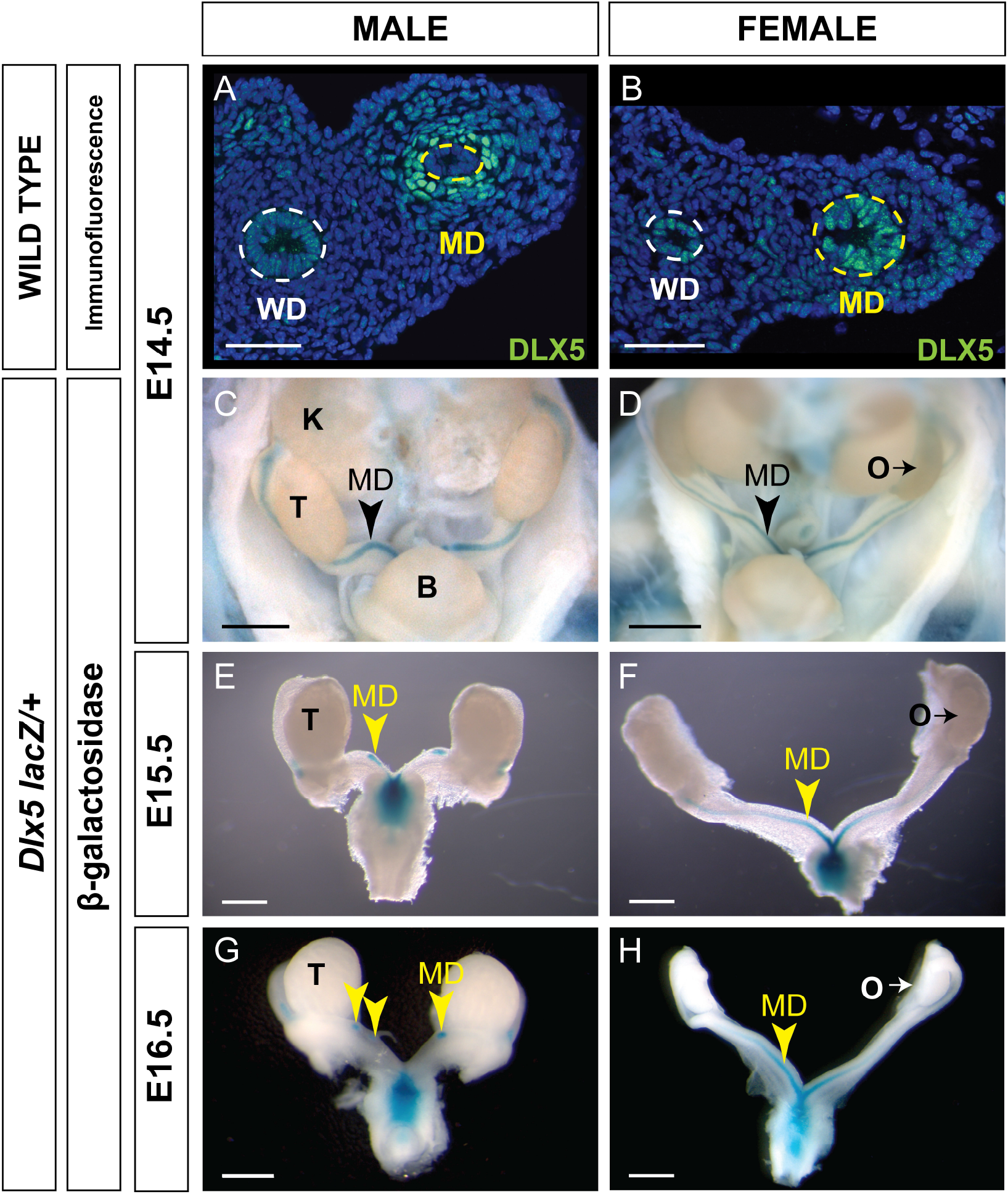
DLX5 and *Dlx5-lacZ* expression in the Müllerian duct. Immunofluorescent staining of DLX5 in cross sections of the mesonephros from E14.5 male (**A**) and female (**B**) embryos. Dotted lines surround the Müllerian duct (MD) and Wolffian duct (WD) epithelium. Scale bars = 50 μm. *Dlx5-lacZ* expression during Müllerian duct development. Wholemount β-gal-stained urogenital organs from E14.5 to 16.5 *Dlx5-lacZ* heterozygous male (**C, E, G**) and female (**D, F, H**) embryos. Arrowheads, Müllerian ducts. B, bladder; K, kidney; O, ovary; T, testis. Scale bars = 500 μm.

We also used a *Dlx5-lacZ* knock-in allele to examine *Dlx5* expression in the developing reproductive organs (19,20). *Dlx5-lacZ* expression was detected at E14.5 in the Müllerian ducts of both male and female embryos (**Fig. 1C, D**). In male and female embryos, β-gal staining was observed throughout the entire length of the Müllerian ducts. At E15.5 and E16.5, *Dlx5-lacZ* expression persisted in the residual Müllerian ducts of male embryos (**Fig. 1E, G**). Consistent with earlier studies, in *Dlx5-lacZ* female embryos at E15.5 and E16.5, β-gal staining was detected throughout the entire length of the Müllerian ducts with strongest staining caudally (**Fig. 1E, G**) (23). These data indicate that *Dlx5* is expressed in the male Müllerian duct mesenchyme and female Müllerian duct epithelium. Thus, *Dlx5* exhibits a sexually dimorphic pattern of expression in the developing Müllerian system.

We used a *Dlx6-lacZ* knock-in allele (20) to examine *Dlx6* expression in the developing reproductive organs (**Fig. 2**). *Dlx6-lacZ* expression in male embryos was detected at E14.5 and E15.5 throughout the entire length of the Müllerian ducts and later at E16.5 as the Müllerian duct was regressing (**Fig. 2A-C**). Histological analysis showed that β-gal expression was localized to the Müllerian duct mesenchyme (**Fig. 2B’**). No β-gal staining was detected in the Müllerian ducts of female embryos at each time point examined (E14.5 to E16.5) (**Fig. 2D-F**). These results show that *Dlx6* is expressed in the male Müllerian duct mesenchyme. The male-specific expression of *Dlx6* in the developing reproductive organs demonstrates that it is expressed in a sexually dimorphic pattern. These findings also show that *Dlx5* and *Dlx6* are co-expressed in the male Müllerian duct mesenchyme at the initiation and during ductal regression.

**Fig. 2.**
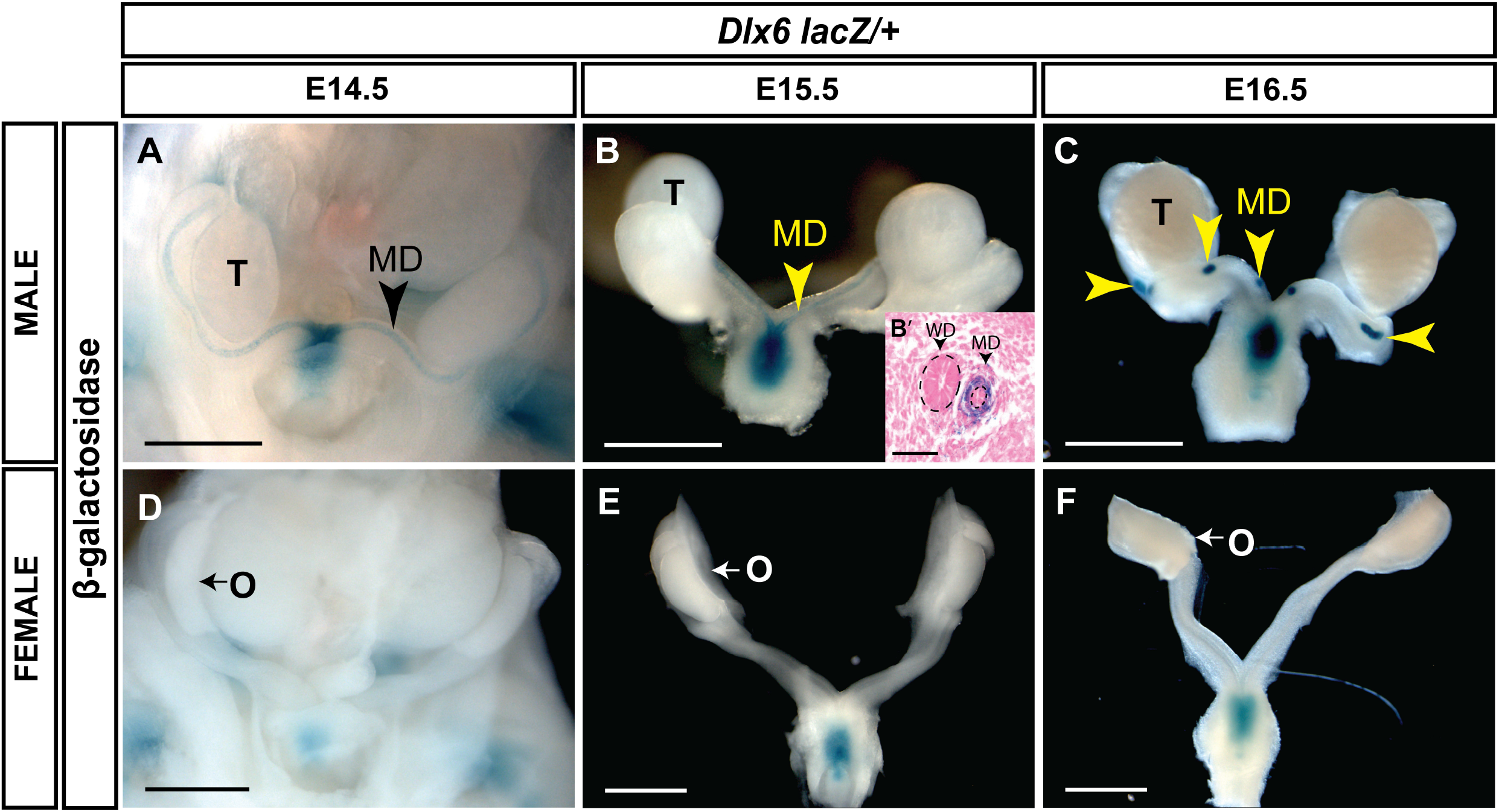
*Dlx6-lacZ* expression during Müllerian duct development. Wholemount β-gal-stained urogenital organs from E14.5 to 16.5 *Dlx6-lacZ* heterozygous male (**A-C**) and female (**D-F**) embryos. Yellow arrowheads, Müllerian ducts. MD, Müllerian ducts; O, ovary; T, testis. Scale bars = 500 μm. Eosin counterstained paraffin section (10 μm) of wholemount β-gal-stained E15.5 *Dlx6-lacZ* heterozygous male (**B’**). Dotted lines surround the Müllerian duct and Wolffian duct epithelium. Arrowheads, Müllerian duct, Wolffian duct. MD, Müllerian duct; WD, Wolffian duct. Scale bars = 50 μm.

### AMH is necessary for *Dlx5* and *Dlx6* expression in the Müllerian duct mesenchyme

The temporal and spatial patterns of expression for *Dlx5* and *Dlx6* in the developing male Müllerian duct mesenchyme are consistent with the idea that they are induced by AMH signaling. To test this idea, we performed genetic experiments examining *Dlx5* and *Dlx6* expression in the absence of AMH signaling in male embryos.

To determine if AMH signaling is required for *Dlx5* expression, we performed DLX5 immunostaining of the Müllerian ducts of *Amhr2 Cre/lacZ* mutant males at E14.5 (**Fig. 3**). *Amhr2-Cre* and *Amhr2-lacZ* are knock-in alleles that are also loss-of-function alleles (17,18). Thus, *Amhr2 Cre/lacZ* mutants lack *Amhr2* function, blocking AMH signaling, resulting in males with a retained and fully developed Müllerian system (8). DLX5 positive Müllerian duct mesenchyme cells were detected in wild-type male controls (**Fig. 3A**). No DLX5 immunostaining was detected in the Müllerian duct mesenchyme of *Amhr2 Cre/lacZ* mutant males (**Fig. 3B**). This suggests that DLX5 expression requires AMH signaling.

**Fig. 3.**
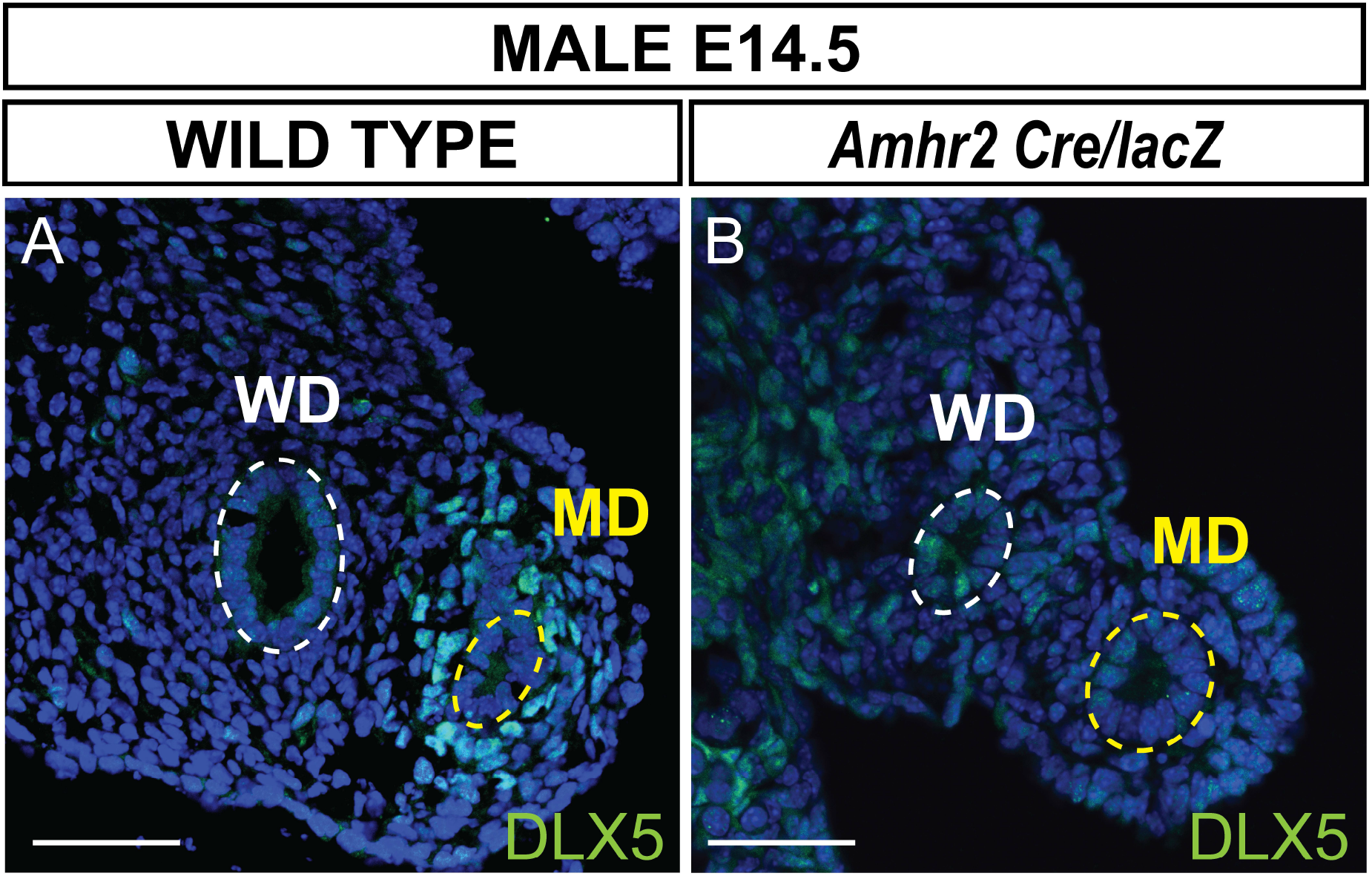
AMH signaling is required for DLX5 expression. Immunofluorescent staining of DLX5 in cross sections of the mesonephros from E14.5 wild type (**A**) and *Dlx5-lacZ/+; Amhr2-Cre/lacZ* (**B**) male embryos. DLX5 is not expressed in *Amhr2-Cre/lacZ* males, indicating AMH signaling is required for DLX5 expression in the male Müllerian duct (MD). Yellow dotted lines surround the MD epithelium; white dotted lines surround the Wolffian duct (WD). MD, Müllerian duct; WD, Wolffian duct. Scale bars = 50 μm.

To determine if AMH signaling was required for *Dlx6* expression, we generated E15.5 *Dlx6-lacZ/+; Amhr2-Cre/Cre* male embryos (**Fig. 4**). β-gal staining was detected in the residual Müllerian duct tissue of *Dlx6-lacZ/+* embryos (**Fig. 4A**). However, no β-gal staining was observed in the Müllerian ducts of E15.5 *Dlx6-lacZ/+; Amhr2-Cre/Cre* male embryos that retained a complete Müllerian system due to the block in AMH signaling **(Fig. 4B**). This suggests that *Dlx6-lacZ* expression requires AMH signaling.

**Fig. 4.**
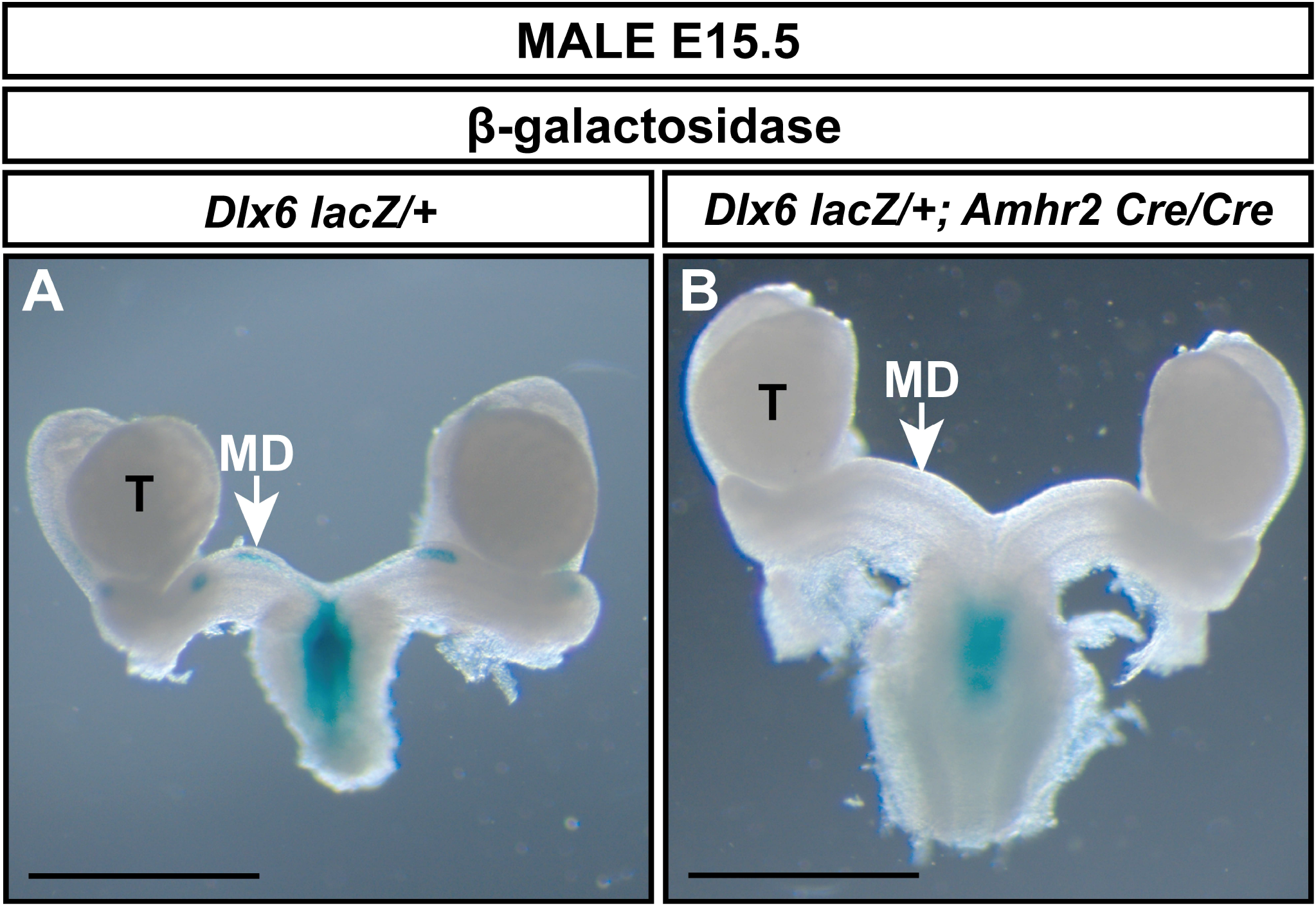
AMH signaling is required for *Dlx6-lacZ* expression in the Müllerian duct. Wholemount β-gal-stained urogenital organs from E15.5 *Dlx6-lacZ/+* (**A**) and *Dlx6-lacZ/+; Amhr2-Cre/Cre* (**B**) male embryos. Arrow in **A**, residual β-gal-stained Müllerian duct (MD) tissue. Arrow in **B**, absence of β-gal-stained Müllerian duct tissue in fully retained Müllerian duct caused by loss of *Amhr2*. MD, Müllerian duct; T, testis. Scale bars=1000 μm.

### *Dlx5* and *Dlx6* regulate Müllerian duct regression

*Dlx5-lacZ* homozygotes die shortly after birth with craniofacial defects (19). In wild-type male embryos, most of the Müllerian duct is regressed at E16.5 and regression is complete by E18.5. We initially screened for the presence of residual Müllerian duct-derived tissues (uterus) by histological analysis of *Dlx5-lacZ* homozygous mutants at E18.5 (**Fig. 5A, B**). None of the 4 *Dlx5-lacZ* homozygous mutant males screened by histological sectioning had uterine tissue (**Fig. 5B**). We next examined Müllerian duct regression in *Dlx5-lacZ* homozygous mutant males and controls at E16.5, using *lacZ* expression to mark the Müllerian duct. Five of 6 *Dlx5-lacZ* homozygous mutant males analyzed had Müllerian duct regression like controls (**Fig. 5C**). However, one of the mutants retained a significant amount of Müllerian duct (**Fig. 5D**). These results indicate that *Dlx5* contributes to Müllerian duct regression.

**Fig. 5.**
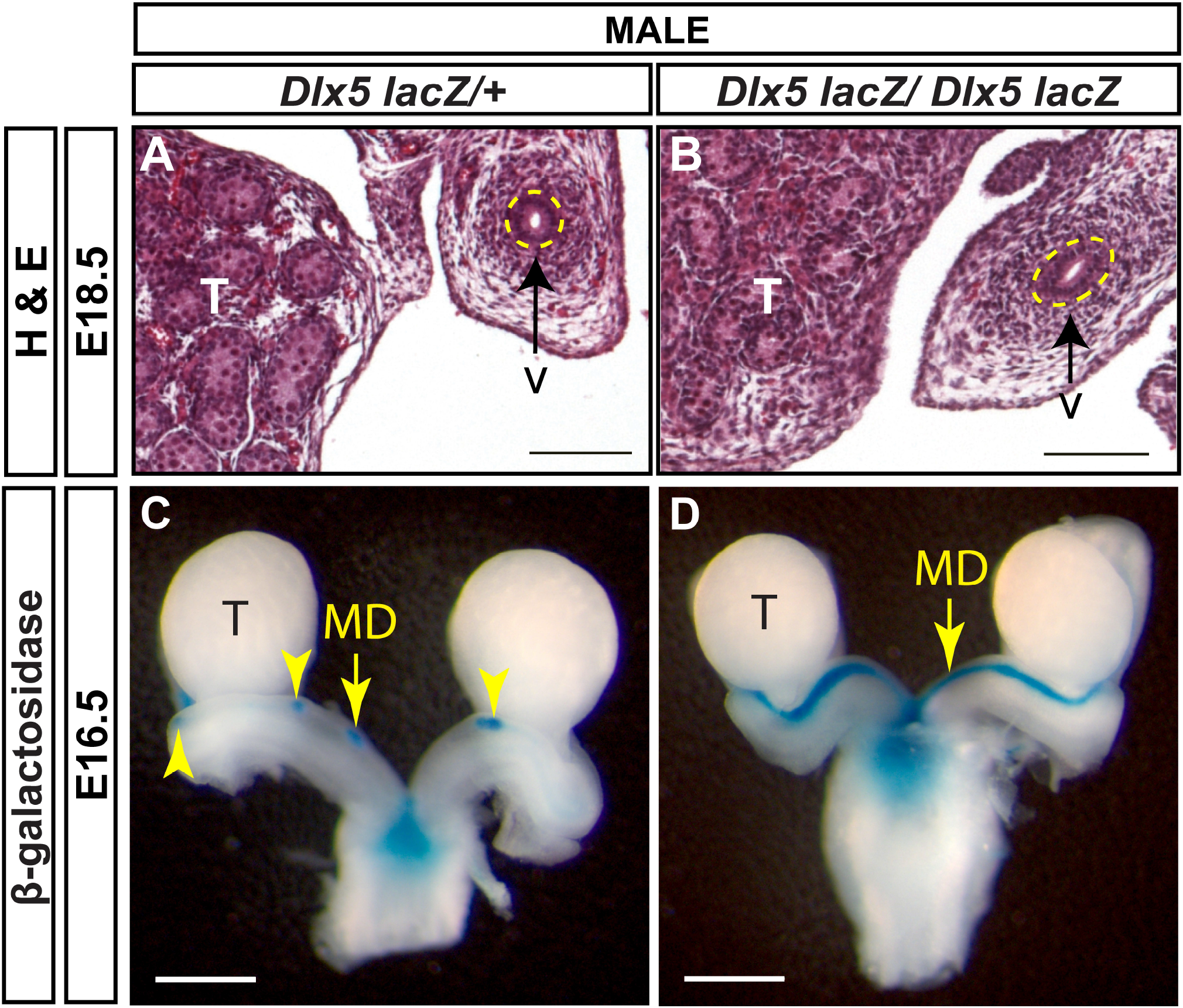
Müllerian duct regression in *Dlx5* mutant males. Hematoxylin (H) and Eosin (E) stained cross sections of the vas deferens (v) adjacent to the testis (T) from E18.5 *Dlx5-lacZ/+* (**A**) and *Dlx5-lacZ/Dlx5-lacZ* (**B**) male embryos. Yellow dotted lines surround the vas deferens epithelium. v, vas deferens; T, testis. Scale = 100 μm. Wholemount β-gal-stained urogenital organs from E16.5 *Dlx5-lacZ/+* (**C**) and *Dlx5-lacZ/Dlx5-lacZ* (**D**) male embryos. Yellow arrowheads, Müllerian ducts. T, testis; MD, Müllerian duct. Scale bars = 500 μm.

*Dlx6-lacZ* homozygous mutants die within a day after birth also with craniofacial abnormalities (20). Like the *Dlx5-lacZ* mutants, we examined Müllerian duct regression by *lacZ* expression in mutants and controls at E16.5 and E18.5. At E16.5, control males showed very small β-gal positive regions, indicating residual Müllerian duct tissue (**Fig. 6A**). In contrast, in E16.5 *Dlx6-lacZ* homozygous mutant male embryos there were still significant stretches of β-gal staining notably in the Müllerian ducts adjacent to the testes (**Fig. 6B**). However, by E18.5, Müllerian duct regression in *Dlx6-lacZ* homozygous mutant male embryos was comparable to control males **(Fig. 6C, D)**. These results suggest that loss of *Dlx6* results in a delay in Müllerian duct regression and *Dlx6* contributes to Müllerian duct regression.

**Fig. 6.**
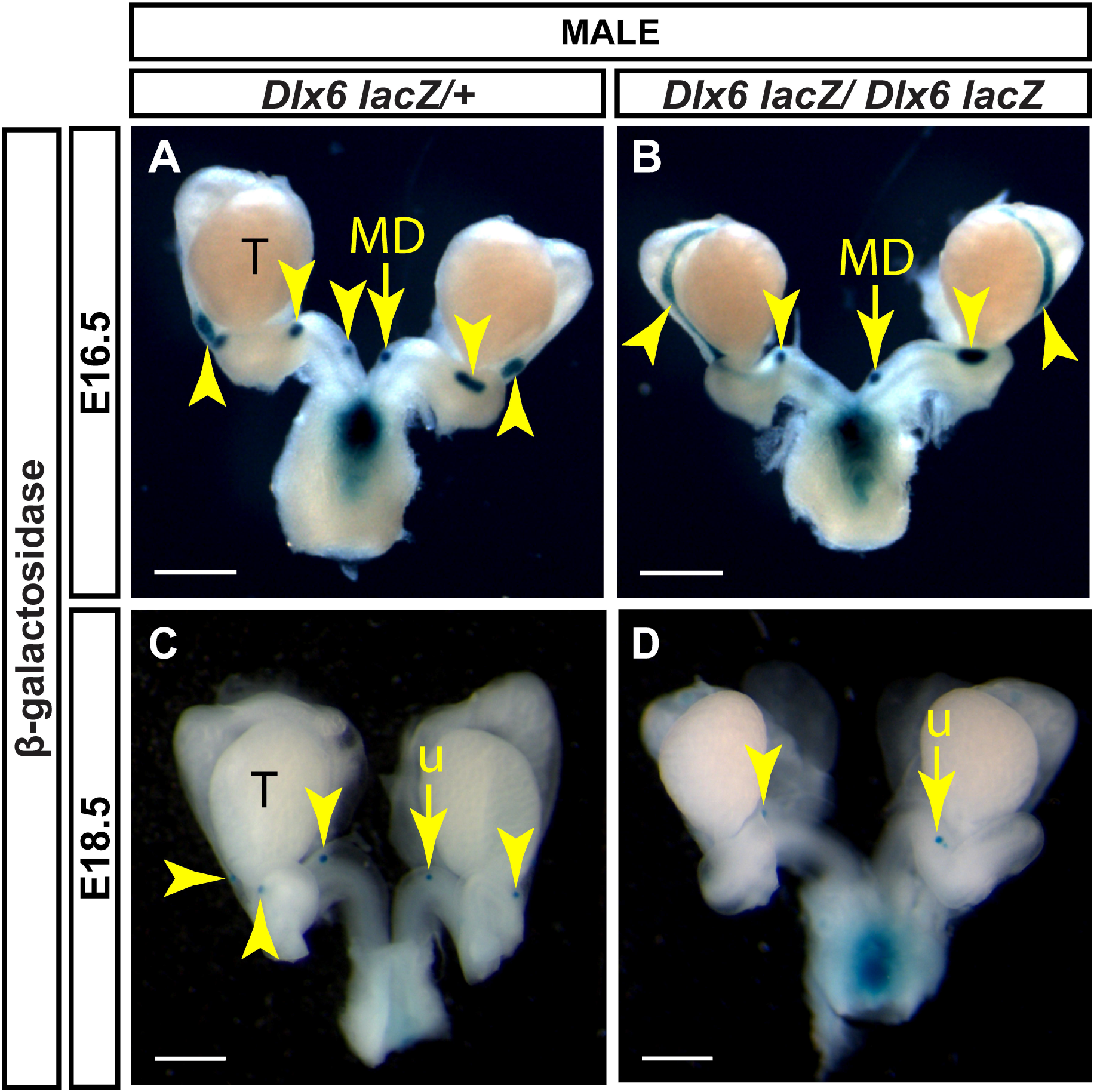
Müllerian duct regression in Dlx6 mutant males. Wholemount β-gal-stained urogenital organs from E16.5 *Dlx6-lacZ/+* (**A**) and *Dlx6-lacZ/Dlx6-lacZ* (**B**) male embryos and E18.5 *Dlx6-lacZ/+* (**C**) and *Dlx6-lacZ/Dlx6-lacZ* (**D**) male embryos. Yellow arrowheads, Müllerian ducts (**A, B**) and uterus (**C, D**). T, testis; MD, Müllerian duct; u, uterus. Scale bars = 500 μm.

Müllerian duct regression in *Dlx5* male homozygous mutants was variable, whereas in *Dlx6* male homozygous mutants it was delayed. *Dlx5* and *Dlx6* are co-expressed in the male Müllerian duct mesenchyme, suggesting that they may act together for Müllerian duct regression. Therefore, we examined Müllerian duct regression in *Dlx5-6* homozygous mutant males. The *Dlx5-6* allele (*Dlx5/Dlx6^tm1Levi^*) is a deletion of both *Dlx5* and *Dlx6* coding regions that also contains a *lacZ* reporter (21). However, the *lacZ* reporter in this allele is not functional. *Pax2* is expressed in the developing Müllerian duct epithelium (24). Therefore, we followed Müllerian duct regression by wholemount immunostaining for PAX2. At E16.5, much like the *Dlx5* male homozygous mutants, one *Dlx5-6* homozygous mutants had near complete retention of the Müllerian duct epithelium while others were comparable to wild type (**Fig. 7A, B**). To quantify the portion of Müllerian duct epithelium retained in *Dlx5-6* homozygous mutants and controls we calculated the ratio of the length of retained Müllerian duct epithelium to Wolffian duct epithelium from the whole mount images. Ratios for *Dlx5-6* homozygous mutants were 0.70 (**Fig. 7B**), 0.32 and 0.27 in comparison to ratios of controls 0.30, 0.25 and 0.19 (**Fig. 7A**). At E18.5 significant portions of uterine epithelium were retained in all *Dlx5-6* homozygous mutants examined in comparison to controls (**Fig. 7C, D**). Ratios of retained uterine epithelium to vas deferens epithelium for *Dlx5-6* homozygous mutants were 0.53, 0.48 and 0.39 (**Fig. 7D**), in comparison to ratios of controls 0.03 (**Fig. 7C**) and 0.04. Consistent with these results, initial histology at E18.5 detected retained uterine tissue in 1 of 4 *Dlx5-6* homozygous mutant males examined (**Fig. 7E, F**). These results suggest that *Dlx5* and *Dlx6* contribute to Müllerian duct regression and may be functioning redundantly.

**Fig. 7.**
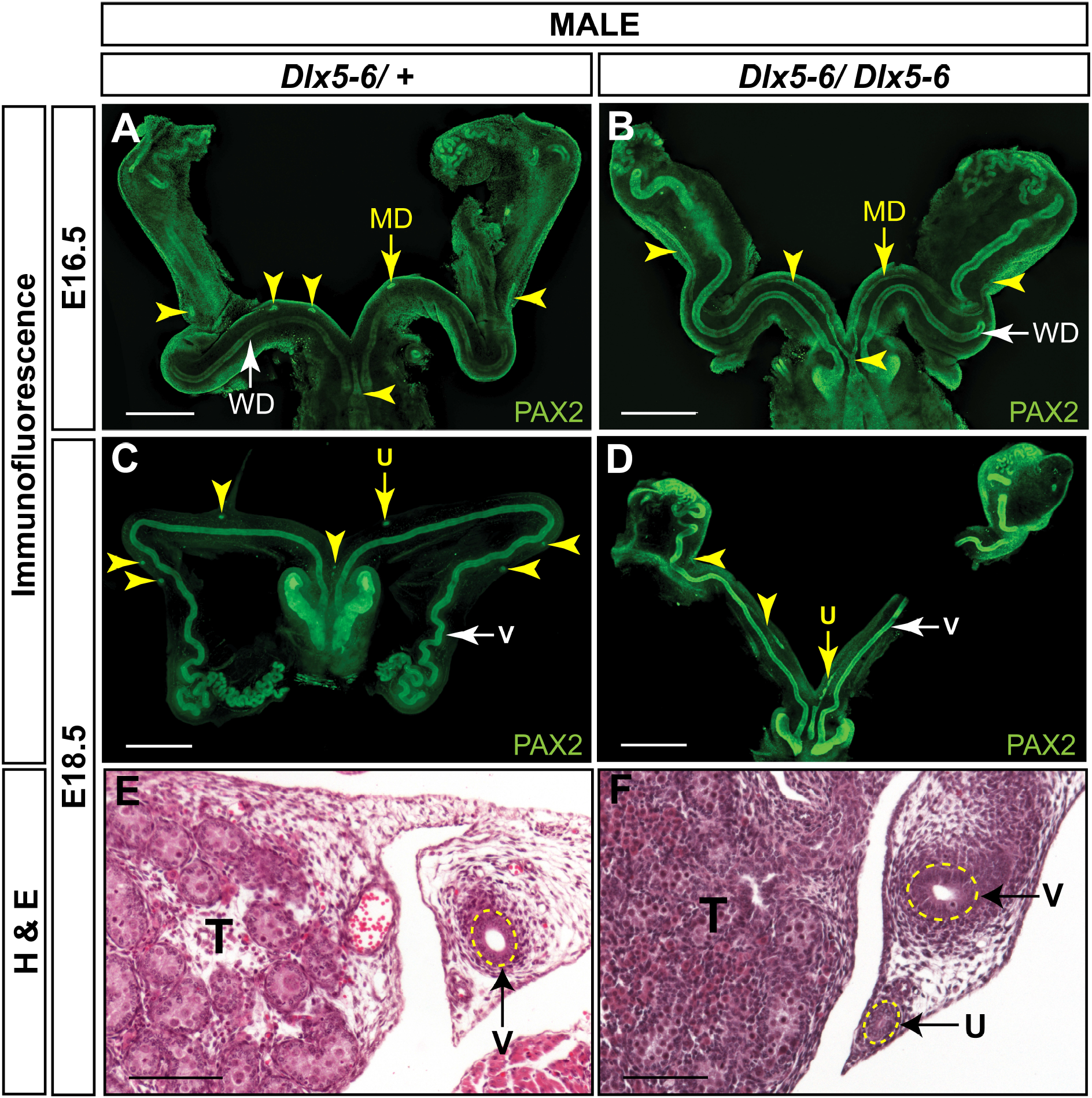
Müllerian duct regression in *Dlx5-6* mutant males. Wholemount PAX2 immunofluorescent staining of E16.5 *Dlx5-6/ +* (**A**) and *Dlx5-6/Dlx5-6* (**B**) male embryos and E18.5 *Dlx5-6/ +* (**C**) and *Dlx5-6/Dlx5-6* (**D**) male embryos. Yellow arrowheads, Müllerian ducts (**A, B**) and uterus (**C, D**). White arrowheads, Wolffian ducts (**A, B**) and vas deferens (**C, D**). MD, Müllerian duct; WD, Wolffian duct; T, testis; v, vas deferens; u, uterus. Scale bars = 500 μm. Hematoxylin (H) and Eosin (E) stained cross sections of the vas deferens (v) adjacent to the testis (T) from E18.5 *Dlx5-6/ +* (**E**) and *Dlx5-6/Dlx5-6* (**F**) male embryos. T, testis; v, vas deferens; u, uterus. Scale bars =100 μm.

## Discussion

Previously, we compared the transcriptomes of FACS-isolated E14.5 male and female Müllerian duct mesenchyme cells in a screen for AMH-induced genes that mediate Müllerian duct regression during male differentiation (16). In our initial analysis, we identified *Osx* as an AMH-induced gene that is expressed in the male Müllerian duct mesenchyme to regulate Müllerian duct regression. Upon further analysis of the transcriptomes, we found that the distal-less homeobox genes *Dlx5* and *Dlx6* were upregulated in males relative to females. We found that both genes are expressed in the male Müllerian duct mesenchyme and are dependent on AMH signaling. The *Dlx5*-null and *Dlx6*-null male embryos (19,20) showed variable retention of the Müllerian ducts and a delay in Müllerian duct regression, whereas the *Dlx5-6*-null male embryos (21) retained more Müllerian duct tissue compared to each single mutant. These findings place a new pair of transcription factors that act together in the gene regulatory network to mediate AMH-induced Müllerian duct regression during male differentiation (25).

### AMH signaling is required for *Dlx5* and *Dlx6* expression in the male Müllerian duct mesenchyme

We found that both *Dlx5* and *Dlx6* are expressed in the male Müllerian duct mesenchyme during the embryonic stages coinciding with the initiation and progression of Müllerian duct regression. *Dlx* genes are organized as linked pairs in the mammalian genome (26). These *Dlx* gene pairs are typically co-expressed in various tissues, suggesting common cis regulation. Perhaps *Dlx5* and *Dlx6* share Müllerian duct mesenchyme-specific regulatory elements that respond to AMH signaling.

Previously, we discovered that *Osx* was expressed in the male but not in female Müllerian duct mesenchyme (16). Our observations of *Dlx5* and *Dlx6* expression add to a growing list of genes that exhibit sexually dimorphic patterns of expression in the Müllerian ducts. We found that the expression of *Dlx5*, and *Dlx6* in the Müllerian duct mesenchyme was dependent upon AMH signaling through AMHR2. Similar results were found for *Osterix* expression (16). *Amhr2* is expressed in the Müllerian duct mesenchyme of both male and female embryos (6,17,18). The female Müllerian duct mesenchyme is competent to respond to AMH because ectopic AMH can induce Müllerian duct regression, eliminating the development of the uterus and oviducts (13). Thus, the sexually dimorphic expression of *Osx*, *Dlx5,* and *Dlx6* expression in the Müllerian duct mesenchyme mirrors the sexually dimorphic expression of fetal AMH.

Interestingly, *Dlx5* is also expressed in the Müllerian duct of female embryos. However, in contrast to mesenchymal expression in male embryos, expression in female embryos was localized to the Müllerian duct epithelium. This epithelial expression persists at later stages of embryogenesis and postnatally in the luminal and glandular epithelium of the uterus (23). We found that *Dlx6* expression in the Müllerian duct was only in male embryos; no expression was detected in female embryos. However, *Dlx6* expression is detected later in the postnatal uterine epithelium (23). These observations suggest that the expression of *Dlx5* in the epithelium of the female Müllerian duct is independent of AMH signaling, suggesting that other factors direct expression in this tissue compartment.

### *Dlx5* and *Dlx6* mediate Müllerian duct regression

Simultaneous deletion of *Dlx5* and *Dlx6* in the postnatal uterus leads to alterations in uterine adenogenesis and infertility, suggesting that they act redundantly for epithelial morphogenesis in the uterus (23). In addition to the postnatal uterus, redundant functions for *Dlx5* and *Dlx6* have also been reported for limb and craniofacial development (19–21,27,28). The overlapping expression patterns of *Dlx5* and *Dlx6* suggested that they functioned redundantly in the male Müllerian duct mesenchyme. Male homozygotes for each single mutation retain variable amounts of Müllerian tissue and male homozygous double mutants retain more Müllerian tissue. Interestingly, the *Dlx5-6* double mutant males did not show complete retention and differentiation of the Müllerian system like *Amh* and *Amhr2* mutant males (7,8). The incompletely penetrant Müllerian duct regression phenotypes in the *Dlx5-6* double mutant male embryos suggests that there are other mediators of AMH-induced Müllerian duct regression that act together with *Dlx5-6* (e.g. *Osx*). Previous studies suggest that there are significantly excess levels of AMH produced for Müllerian duct regression because *Amh* levels must be reduced to ~10% of wild type to uncover partial Müllerian duct regression phenotypes (18). Perhaps downstream target genes of AMH signaling must be collectively inactivated for a complete block to Müllerian duct regression (29,30).

*Dlx* genes are known targets of BMP signaling (26). Interestingly, AMH signal transduction shares receptors (BMPR1A and ACVR1) and downstream R-SMAD proteins (SMAD1, 5, 8) with BMP signaling (Jamin et al., 2002; Orvis et al., 2008). *Osx* was discovered as a BMP-induced gene (15). *Osx*, *Dlx5,* and *Dlx6* each have roles in skeleton development and Müllerian duct regression (15,16,19–21,27,28). Do these transcription factor genes regulate the same downstream effectors in skeleton-forming and Müllerian duct mesenchyme cells, or do they regulate different sets of target genes? This question can be addressed by identifying tissue specific cis regulatory elements that are bound by OSX, DLX5, and DLX6 and regulate transcription.

## Author Contributions

R.D.M. and R.R.B. conceived the study and designed experiments. R.D.M. performed experiments and analyzed data. B.B. and G.L. generated mutant embryos. R.D.M. and R.R.B. wrote the paper with input from all authors.

## Acknowledgements

We thank Dr. Ying Wang for advice on immunofluorescent studies, Kazi Hossain for assisting with imaging and the members of our labs for helpful discussions.

